# Phenotype Prediction using a Tensor Representation and Deep Learning from Data Independent Acquisition Mass Spectrometry

**DOI:** 10.1101/2020.03.05.978635

**Authors:** Fangfei Zhang, Shaoyang Yu, Lirong Wu, Zelin Zang, Xiao Yi, Jiang Zhu, Cong Lu, Ping Sun, Yaoting Sun, Sathiyamoorthy Selvarajan, Lirong Chen, Xiaodong Teng, Yongfu Zhao, Guangzhi Wang, Junhong Xiao, Shiang Huang, Oi Lian Kon, N. Gopalakrishna Iyer, Stan Z. Li, Zhongzhi Luan, Tiannan Guo

## Abstract

A novel approach for phenotype prediction is developed for mass spectrometric data. First, the data-independent acquisition (DIA) mass spectrometric data is converted into a novel file format called “DIA tensor” (DIAT) which contains all the peptide precursors and fragments information and can be used for convenient DIA visualization. The DIAT format is fed directly into a deep neural network to predict phenotypes without the need to identify peptides or proteins. We applied this strategy to a collection of 102 hepatocellular carcinoma samples and achieved an accuracy of 96.8% in classifying malignant from benign samples. We further applied refined model to 492 samples of thyroid nodules to predict thyroid cancer; and achieved a predictive accuracy of 91.7% in an independent cohort of 216 test samples. In conclusion, DIA tensor enables facile 2D visualization of DIA proteomics data as well as being a new approach for phenotype prediction directly from DIA-MS data.

## Introduction

Phenotype prediction is a central task for biomedical research and clinical decision making, which is typically dependent on phenotypic measurement, such as blood pressure monitoring [1], electrocardiogram [2] at the macro level as well as high throughput omics sequencing technology of the genome [3], DNA methylome [4], mRNA [5], proteins [6] and metabolites [7] at the molecular level. Mass spectrometry (MS)-based proteomics [8] has demonstrated its ability to measure thousands of proteins in complex biological samples within several hours using data dependent acquisition (DDA) MS and data-independent acquisition (DIA) MS. In DIA-MS, exemplified by SWATH [9], all flyable peptide precursors are recursively fragmented in parallel in isolated m/z windows, and all the fragment data, together with the peptide precursor data, are comprehensively recorded. The resultant DIA data therefore serve as a permanent digital map representing all the measurable protein signals that is re-minable to test the new hypothesized biomarkers [10]. Emerging high-throughput sample preparation methods such as pressure-cycling technology (PCT) offer rapid and reproducible processing of minute amounts of biopsied tissue archived as formalin-fixed paraffin-embedded (FFPE) samples [11] which can be then proteome-digitized by DIA-MS as a permanent digital archive [12].

DIA maps are usually analyzed with pattern match algorithms such as OpenSWATH [13]. In DIA data, co-eluting precursors may be co-fragmented in the same window which generates highly convoluted fragment spectra. This complexity can be deconvoluted using prior information such as a highly curated spectral library containing peptide precursors of interest, which includes precursors’ m/z and the m/z of their fragment ions with corresponding relative intensities and calibrated retention times (RT). The software performs targeted extraction of ion chromatograms (XIC) from DIA maps for selected peptide precursors based on mzXML [14] or mzML format [15]. Together with other features, such as errors in mass and RT, and peak shape, a discriminating statistical learning model is built to score the true target based on target-decoy approach for peptide identification [13]. Several deep learning-based algorithms have been developed recently to directly predict spectral library and to reduce bias from DDA experiments [16]. Alternatively, the DIA data can be analyzed by spectrum-centric library-free approach [17] which determines precursor-fragment pairs from precursor ions and fragment ions, uses theoretical fragment ion predictions to query and score the multiplexed MS2 spectra of SWATH-MS data. These tools will usually give rise to a distinct identified and quantified peptide or protein matrix which are subject to downstream statistical learning procedures for classification of disease types and feature selection of biomarker candidates [18]. However, these tools analyze only a small portion of the DIA map, leaving many biological signals uninterpreted. Erroneous estimation of protein intensities and technical missing values are additional challenges for effective statistical analysis, including deep learning.

Deep learning, or deep neural networks (DNN), learns from training data to extract effective features and performs classification tasks in an optimal way [19]. Instead of relying on handcrafted assumptions and knowledge, the deep learning methodology learns domain knowledge from data and delivers the best models that fit reality. It has surpassed human performance in image classification on the ImageNet dataset [20] as Large Scale Visual Recognition Challenge. In particular, ResNet [21], a special type of convolutional neural network (CNN), has demonstrated advantages over conventional CNN. It has been widely used for object detection and classification in computer vision.

In this work, we propose a novel MS data analysis approach for phenotype prediction. First, a novel data format called “DIA tensor” (DIAT) is proposed as a preprocessing step and also as a convenient means of data visualization. Second, the pooled DIAT map is divided into blocks and fed into a deep neural network, a modified version of ResNet-18 to predict the phenotype probabilities for benign vs. malignant classification of clinical samples. Finally, the prediction scores of all blocks are fused to make the final prediction. This approach works directly on MS data, without the need to identify peptides or proteins, avoiding the computationally intensive and error-prone XIC procedure.

To prove the concept of our approach, we performed the classification task on a set of 102 human liver tissue samples consisting of benign liver and hepatocellular carcinomas. We further applied these procedures to a more challenging clinical problem of diagnosing the malignant vs. benign status of thyroid nodules. We showed that deep learning using the pooled DIAT data format could provide an alternative approach for MS-based clinical diagnosis.

## Results

### DIAT enables end-to-end phenotype prediction without peptide identification

Here we developed DIA tensor (DIAT) as a new format to represent DIA-MS data that can be used directly for end-to-end deep learning classification model, skipping the peptide identification step by the peptide centric [13] or spectrum centric [17] approach of DIA MS data. The DIAT is a three-dimensional tensor to represent the pooled MS2 ion maps, consisting of MS1 precursor window index, cycle index, and pooled fragment m/z. To transform a DIA raw file into DIAT format, the vendor mass spectrometry DIA raw files were first converted into mzXML files by MSconvert [22]. Key attributes such as the scan level, scan index, precursor center m/z, fragment m/z and their intensities were extracted from the open data format mzXML files. The mzXML file of DIA-MS data is recorded by consecutive repeated cycles throughout the LC time range. Each cycle contains one MS1 spectrum and the corresponding fragment ion MS2 spectra stepping through a sequential discrete precursor isolation window (**Figure 1A**). Some scans in a cycle may be missing occasionally due to MS analyzer instrument error, which will cause the subsequent scan sequence numbers to be misaligned. To keep the scans in order, the missing scans were detected and filled with zero values. The m/z values were binned with the size of the mass accuracy of the mass analyzer. The MS2 fragment ion spectra of the same precursor window are aligned together accordingly by scans tensor generation, similar to the concept of extracted ion chromatograph [9] to depict the elution pattern of a peptide precursor or its fragment ions. We then generated a three-dimensional tensor of reordered MS2 intensity as binned m/z reordered cycle index, and precursor window index as DIA tensor. (**Figure 1C**).

**Figure 1.**
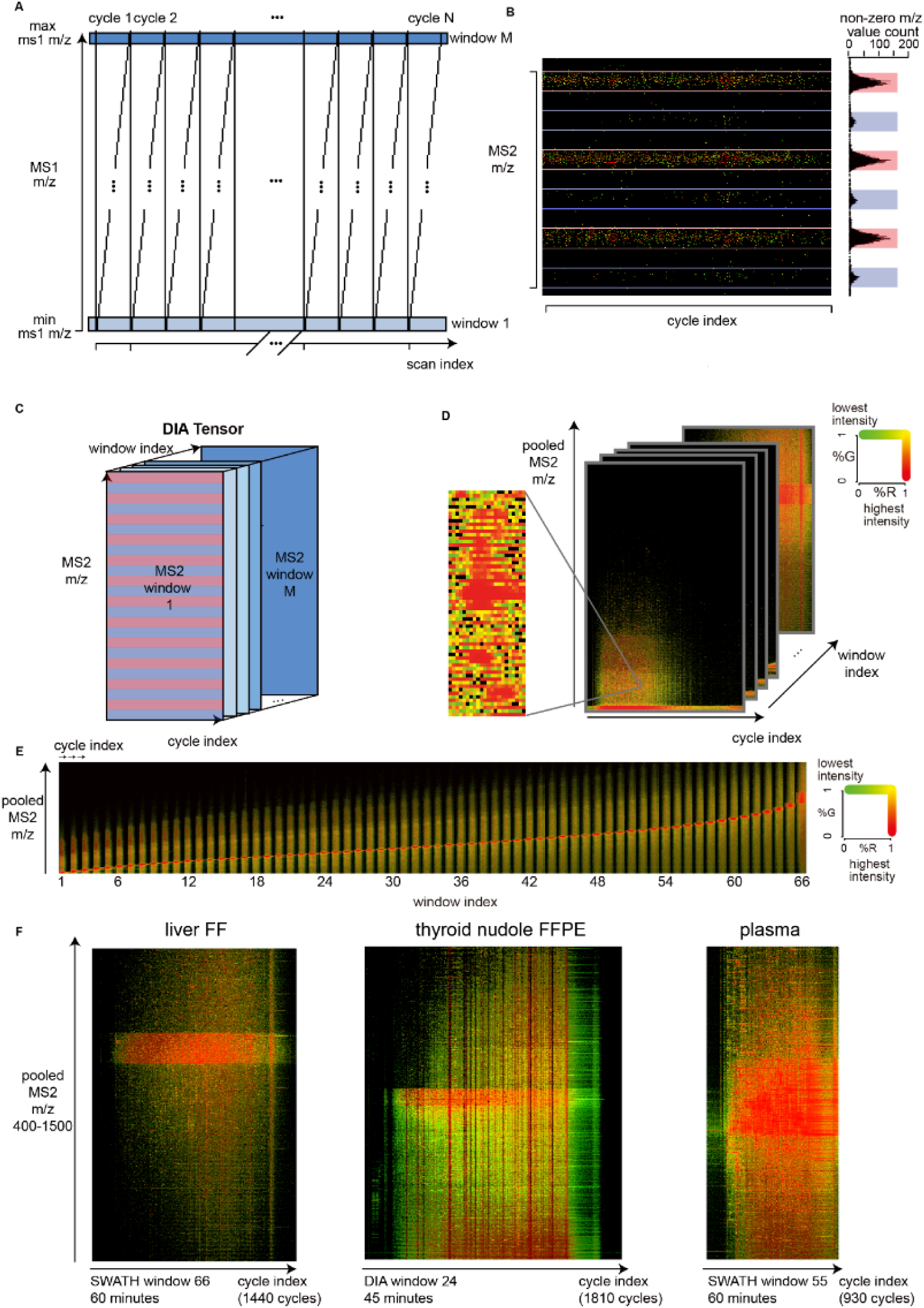
Establishment of DIA tensor. A) Illustration of raw data structure of DIA-MS acquisition scheme, which is a two-dimensional array of MS2 m/z and ion intensities arranged by sequential scans of repeated cycles of one MS1 scan plus a series of MS2 scans that span the entire MS1 precursor m/z range. B) Left panel, image crop of the raw MS2 intensity shows fragment ion clusters in rows. Right panel, the nonzero intensity count distribution of fragment ion m/z from an empirical DIA map. Major peaks are colored in red, while the minor peaks are in blue. C) Layout of the intensity tensor is illustrated by pooled MS2 m/z with reordered scans by cycle index and precursor window index. D) Visualization of DIA tensor. An orthogonal linear sampling was used to sample the Red-Green space to generate a color scheme of green (lowest)-to-red (highest). Black is used for zero values. Zoomed in is a sample visualization in the first isolation window. E) A sample image visualization by flattening the window index dimension. The MS2 windows were sorted by precursor centers accordingly, where the continuous clear red line indicates the corresponding precursor windows of unfragmented precursor ions. F) Visualization of the last window of three different types of clinical samples with different DIA acquisition scheme and gradient length. Liver fresh frozen: SWATH 66 windows, 60 minutes; Thyroid nodule FFPE: DIA 24 windows, 45 minutes; Plasma: 55 windows, 30 minutes.

The size of original DIA tensor is nevertheless too large to feed into the deep learning module and for visualization. Interestingly, we found that the combinations of sequential amino acid m/z values are aggregated into discrete values, which we consistently observed as a pattern of alternating peaks (**Figure 1B**) with predefined grids in the MS2 m/z dimension when counting the nonzero intensity. We further confirmed this pattern by simulation of single charged fragment m/z of human proteome where we also found the same pattern of major peaks (**Figure 1B**) and minor peaks which could be interpreted as doubly charged fragment ions. We then utilized this feature to further reduce the tensor in the m/z dimension. To dynamically determine the pooling size of different m/z ranges, nonlinear squared Gaussian fit of non-zero m/z peaks was used to determine the pooling boundary. By dropping all the grid without peaks and binning major/minor peak area to one row, we reduced the m/z dimension by fifty-fold with the m/z intensity pooled as the sum of the intensities within the pooling boundaries. To visualize the tensor (**Figure 1D**), we discretized the non-zero intensity space and assigned three channel RGB colors, sampled from green to red bidirectional color space, and flattened 2D images by window index, as shown in representative samples (Figure 1E, 1F). The stellar DIA image depicts the MS2 windows sorted by precursor centers, where the continuous clear red line indicates the corresponding precursor windows of unfragmented precursor ions. For deep learning, we used a single channel grey color to match the discretized value and avoided introducing extra redundancy in color dimension. The file size of pooled DIAT is 10-fold smaller than mzXML by SWATH and 2-fold smaller than mzXML by DIA.

### Deep learning based on DIAT using a hepatocellular carcinoma cohort

Modified neural networks of ResNet-18 (see Materials and Methods) was applied to DIAT data directly for phenotype prediction in cancer diagnosis, without relying on the computationally intensive and error-prone XIC procedure. Each tensor was divided into sub-tensors as multi-channel images with a size of pixel 224×224×C (**Figure 2A**) where 224 is the size of the input layer used in ResNet and C is the number of precursor isolation windows. To fit the full size of the tensor, some of the regions on the boundary have partial overlap.

**Figure 2.**
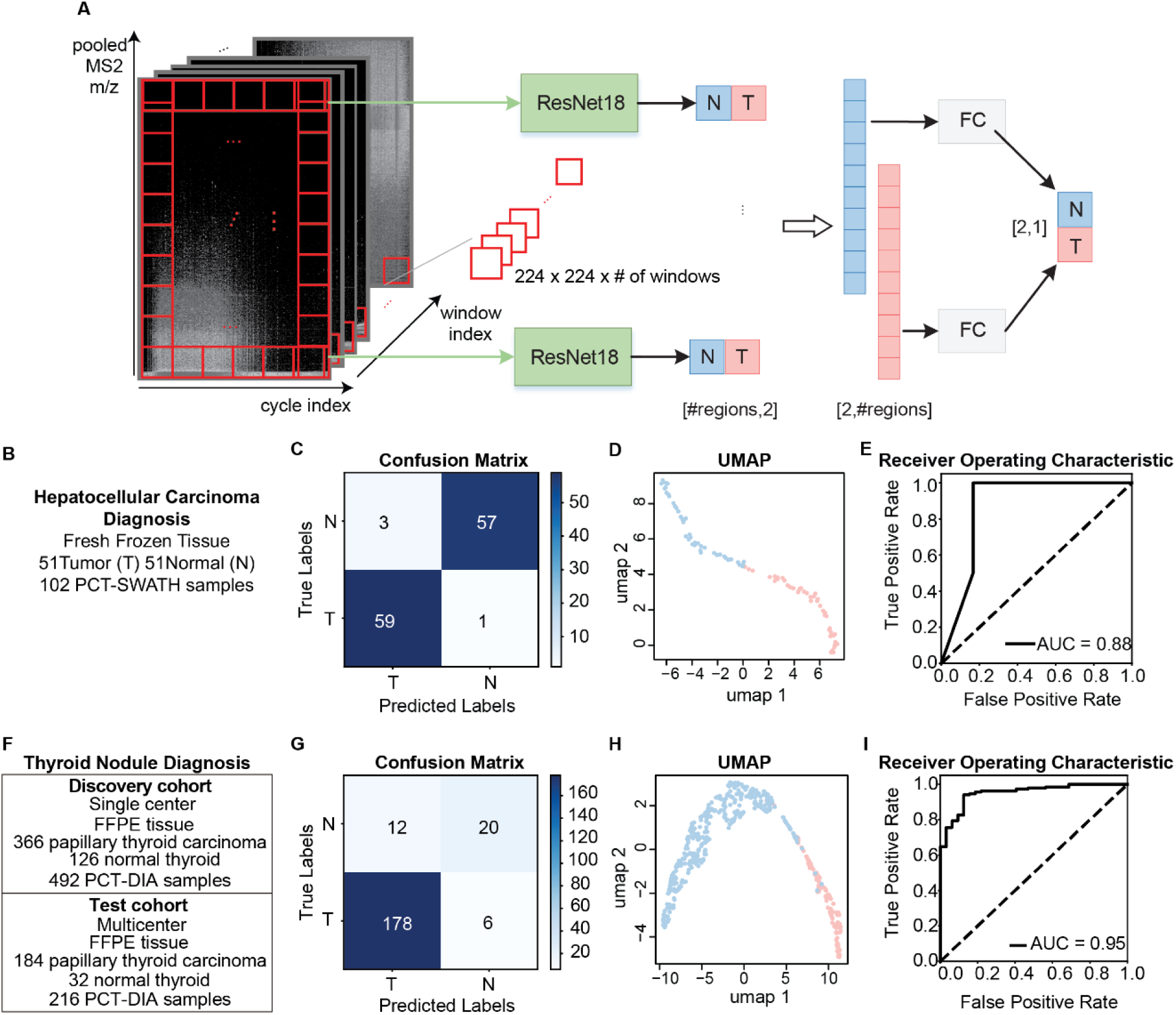
DIA tensor coupled with deep learning for phenotype prediction. A) Overview of a general deep learning framework. DIA tensor was divided into sub tensors of 224 x 224 x number of isolation windows, followed by modified ResNet-18 analysis on each region, and fusion of the results of each region through two fully connected layers. B) The design of HCC PCT-SWATH experiment. C) The total confusion matrix of 10 training experiments. D) The UMAP of training and validation sets of HCC samples E) The ROC curve of the HCC experiment of one test set. The AUC is 0.88. F) The design of thyroid nodule PCT-DIA experiment. G) The confusion matrix of the thyroid nodule diagnosing experiment. H) The UMAP of training cohort of thyroid nodule samples I) The ROC curve for diagnosing malignant thyroid nodules.

A modified ResNet-18 network was applied to each region to obtain prediction scores and the prediction scores of all regions were fused to get the final prediction through a fusion module. The structure of the modified ResNet-18 was the same as conventional ResNet 18 except that the output size of the last fully connected layer was modified to suit the binary classification problem. From this, the output of all regions formed two vectors according to the node type (benign vs. malignant). Finally, two fully connected layers were used in the fusion module to perform the weighted average on two vectors, the output being the final prediction. (**Figure 2A**) A cohort of fresh frozen tissue biopsies of primary liver cancer, hepatocellular carcinoma (HCC) [23], was used in this study. The data set used here is described in greater details in a separate manuscript in preparation. For each patient, tumor and non-tumorous tissue samples were collected and processed with the PCT-SWATH workflow [24] to generate 102 SWATH MS files. We used 70 sample files as the training set and 20 sample files as the validation set, and 12 sample files as the test set. Since there was insufficient data in the test set, ten trials of training were performed, attaining a mean accuracy of 96.8%. (**Figure 2B**) The accuracy was 96.8%, the precision 95.2%, the recall 98.3%, and F1-score 0.967 (**Figure 2C-D**). The hyperparameters such as learning rate, number of iterations, batch size, and loss function were optimized in the HCC dataset so that we could use for the other proteomic data sets.

### Application on cohorts of papillary thyroid carcinomas

To prove that our methodology is generic, we further applied it to a different DIA data set of thyroid nodules which is bigger and more complicated. We adopted the same framework and hyperparameters as in the HCC experiment. Thyroid nodules are a common endocrine disease affecting approximately 50% of the population globally, particularly in women [25]. Nevertheless, only 7-15% of them are malignant and papillary thyroid carcinoma (PTC) accounts for 85% of malignant nodules [26]. Patients suffering from clinically indeterminate nodules are often advised to have surgical resection. However, most resected nodules turn out to be benign after histopathological evaluation, leading to pervasive overtreatment in many countries. PTC has one of the lowest mutation densities compared with other cancers [27], and so is not readily detected by genomic technology. Therefore, we applied PCT-DIA to acquire a total of 708 thyroid DIA files consisting of 158 normal thyroid DIA files (N) and 550 papillary thyroid carcinoma DIA files (P) [28]. The discovery cohort was collected from single hospital and the test cohort was from three different hospitals (**Figure 2E**). The discovery cohort comprised 126 N and 366 P DIA files; the test cohort consisted of 216 DIA files. The data set used here is described in greater details in a separate manuscript under review. Applying the same framework and hyperparameters as in the HCC experiment, we attained an accuracy of 91.7%, precision of 94.6%, recall of 95.7%, and F1-score of 0.951 (**Figure 2F-G**) at the sample level. These results show that our methodology is generic and can be potentially applied to address a broader range of clinical questions.

## Discussion

Although a plethora of mass spectrometric methods enables generation of precise mass spectral data in high-throughput and large volume, application of these data to (clinical) phenotype prediction are mostly performed after molecular interpretation of the mass spectral data [8]. Direct use of mass spectrometric data is used in mass spectrometry imaging technique, often referred to as Matrix-Assisted Laser Desorption/Ionization–Mass Spectrometry Imaging (MALDI-MSI) [29, 30], although ionization methods other than MALDI have been developed too [31]. Currently this field is mainly focused on generating high spatial resolution spectral data for maximal number of molecules, and correlation with histology images [32, 33]. The throughput to generate an image of high spatial resolution is still low. Moreover, the images thus generated can be hardly be the source for deep learning models for phenotype prediction. In addition, clinical application of MALDI-MSI is constrained to analyzing biopsies tissues but is incapable of measuring other types of clinical samples such as unsectioned fresh frozen tissue, blood plasma or serum and stool.

DIA-MS data can be visualized [34, 35] by manual inspection of DIA ion maps, but such visualization is not designed as the input for deep learning model where the large input of m/z data by high-resolution mass analyzers needs to be properly handled. In this work, we have established an end-to-end methodology allowing direct deep learning analysis of DIA-MS raw data in the format of DIAT for phenotype prediction. We first applied this method to distinguish between benign and malignant liver tissues as proof of principle. Then the same method was deployed to predict malignant thyroid nodules. Our method bridges DIA-based proteomics directly to deep learning technology such as residual neural network for computer vision. With this new approach, DIA data do not need to be processed through heavy computation for interpreting the mass spectra data, thereby circumventing errors and distortions artificially introduced during mass spectral interpretation. As the quality and quantity of MS data increase, we can foresee the potential of DIAT in rapid clinical diagnosis. Future research is required to interpret the biological insights revealed from deep learning of the DIAT maps. Our preliminary data show this data format and data analysis strategy also applies to DIA data of metabolomes and lipidomes.

## Acknowledgements

This work was supported by National Natural Science Foundation of China (General Program) (Grant No. 81972492 to T.G.), Zhejiang Provincial Natural Science Foundation for Distinguished Young Scholars (Grant No. LR19C050001 to T.G.), Hangzhou Agriculture and Society Advancement Program (Grant No. 20190101A04 to T.G.). We thank Mr. Guan Ruan for help in graphics.

## Authors contributions

T.G., S.Z.L, Z.L., F.Z. designed the project. F.Z., S.Y., L.W. and Z.Z. developed the algorithms and programs with guidance from T.G., S.Z.L. and Z.L.. Y.S. generated the thyroid data with samples procured by S.S., L.C., X.T., Y.Z., G.W., Y.X., OLK., and N.G.I.; X.Y. generated the liver data with samples procured by J.Z., C.L., P.S. and S.H.. F.Z. and T.G. wrote the manuscript with inputs from all co-authors. T.G., S.Z.L. and Z.L. supported and supervised the project.

## Materials and Methods

### Patients and tissue samples of HCC

All tissue samples of hepatocellular carcinoma cohort were collected from Union hospital, Tongji Medical College, Huazhong university of Science and Technology, Wuhan, China. Tissue samples were collected within 1.5 hour after hepatectomy, then frozen and stored at −80°C or temporally in −20°C before moving to −80°C. For each patient, two tissue biopsy punches (with dimension of 5 x 5 x 5mm) including a tumorous tissue and a non-tumorous tissue from an adjacent region as determined by histomorphology.

### PCT lysis and extraction of proteins

About 1mg of the frozen tissue on average was lysed with the PCT-Micro Pestle device in 30μL of lysis buffer composed of 6M urea, 2M thiourea and 0.1M ammonium bicarbonate in a barocycler HUB440 (Pressure BioSciences Inc, South Easton, MA, USA), where the tissue lysis was performed under a program consisting of 60 oscillating cycles, where each cycle consists of high pressure at 45,000psi and 10s of ambient pressure at 30°C. Then the extracted proteins were reduced and alkylated by incubation with 10mM tris(2-carboxyethyl) phosphine (TCEP) and 40mM iodoacetamide (IAA) under gentle vortexing at 600 rpm for 30min (ambient pressure) at 25°C in the dark. Afterwards, proteins was first digested with lys-C (enzyme-to-substrate ratio, 1:40) in the barocycler under the program of 45 cycles of 50s high pressure at 20,000psi 10s ambient pressure, followed with trypsin (enzyme-to-substrate ratio, 1:50) of 90 cycles of at 50s at pressure 20,000psi and 10s ambient pressure. After digestion, the peptides were acidified with trifluoroacetic acid (TFA) to pH 2-3 and cleaned with The Nest Group C18 (17-170ug capacity) and dried under vacuum. Peptides were reconstructed in HPLC-grade water containing 0.1% formic acid and 2% acetonitrile before mass spectrometry injection.

### Mass spectrometry of HCC samples

One microgram of peptide sample was injected to an Eksigent 1D+ Nano LC systems (Eksigent, Dublin) and analyzed in a 5600 TripleTOF mass spectrometer (SCIEX) in SWATH mode. The LC gradient was reduced to 60 min, and the SWATH acquisition scheme to 66 variable windows (Table S1). The other SWATH parameters were set exactly as in our previous study except that the ion accumulation time for each SWATH window was 40 ms. Ion accumulation time for peptide precursors was set at 50 ms. The 102 samples were injected into the MS in randomized sequence once and then the same sequence was injected again to obtain a duplicate. After each gradient, the column was washed twice using ramping gradient to minimize carryover. Mass calibration using beta-gal was performed every fourth injections.

### Dewaxing, rehydration and hydrolysis of FFPE tissues

For each case, three biological replicates of FFPE punches were processed. Before dewaxing, specimens were weighted and recorded. FFPE tissue samples were washed with heptane (Sigma) and 100% ethanol (Sigma), 90% ethanol, 75% ethanol successively in room temperature. After dewaxing and rehydration, 0.1% formic acid (Sigma) was added for achieve C-O hydrolysis of protein methylol products and then washed with 100 mM Tris-HCl (pH=10, Sigma) to exchange the condition for following reaction. Base hydrolysis was performed under 95 degree centigrade using 100 mM Tris-HCl (pH=10). Finally, cool down immediately at 4 Celsius degrees.

### Patients and tissue samples of thyroid nodules

The samples included in our study were FFPE punches collected from four clinical centers from Singapore and China with the approval of the ethics in the indicated hospitals.

In the discovery dataset, FFPE specimens from patients with thyroid adenomas, nodular goiters or different thyroid cancer types, whom were treated in the Singapore General Hospital (SGH) between the year 2012-2017, have been retrospectively retrieved from the Pathology Department of SGH. Haematoxylin and eosin-stained slides from tissue blocks of every single patient were carefully reviewed by an experienced histopathologist who marked out the disease region for tissue coring.

The selection of cases was excluded with extensive thyroiditis and/or inflammation and the pathological lesions of the cases were more than 1 cm in diameter. Pathologists viewed and marked haematoxylin-eosin stained slides and then each case were made punch cores at the region of interest using a hollow metal needle of 1 mm internal diameter and 1 mm length. Each dry FFPE punch weight around 0.6 mg including wax.

### mzXML file converting options

The .wiff and .raw files were converted to mzXML files using MSconvertGUI (64-bit Version: 3.0.19172-57d620127) with parameters peakPicking set as vendor msLevel=1-2

### DIA tensor data structure

Data type: uint16 / uint32 (automatically selected according to the needs of stored data)

Data space: 4D

Z dimension: ms2 window index

Y dimension: m/z index in 0.01 Da bin

X dimension: cycle index (requires cycle alignment for the same batch of datasets)

Color dimension: ms2 intensity (saved as integer type, without any other processing)

The difference of raw DIAT and pooled DIAT:

1. The m/z index changed from the bin number of 0.01 Da bin to the m/z main and sub-peak division area, and then the main and sub-peak numbers after 95% confidence interval fitting.
2. MS2 intensity becomes integer type value after m/z main and auxiliary peak pool

### Binning of m/z

Binning indices: 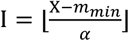 Binning value: *z*_*k*_ = Σ_k_(*Y*|*I* = *k*), *k=*1,…,*N*; where *m*_*min*_ = 400, *m*_*max*_*=* 1500; *α:* bin size 0.01; *X: m/Z* values for a scan; *I:* Indices after binning for a scan; *Y:* Intensities for a scan; *N* = (*m*_*max*_ − *m*_*min*_)/*α z*_*k*_:Binned intensity in bin *k*; if there is no value in bin *k, Z*_*k*_ is 0.

### Pooling of m/z

We count the number of m/z values in a bin to obtain the count distribution of m/z. We assume there is a repeating pattern in the m/z distribution (**Figure S4A**): major peak, no peak, minor peak and no peak with a fixed grid size (0.250132) and peak position, so that the peak apex can be designated into a grid. The original m/z range from 400 to 1500 will be transformed into 2199 grid lines (remove no-peak lines). For grid line with major and minor peaks, the peak center, peak standard deviation, and peak height are fitted by the gaussian function using the 25 points around the peak apex in the pre-designated grid. The pixel value for this grid line is determined by the sum of all the intensities within 1.96 standard deviation by lines. If we are not able to fit a proper peak, it will use the default args (sigma=0.03) to sum all the intensities within 1.96 standard deviation (95% confidence) by lines (**Figure S4B**).

### Simulation of the single-charged human proteome

The human proteome fasta file was downloaded from Uniprot (n2019 August). Monoisotopic mass of amino acid is downloaded from Unimod.org. The proteins are firstly trypsinized *in silico* into peptides with K/R terminus. Unique peptide sequences with length of six to thirty are retained. The fragment mass-to-charge ratio is binned with size 0.01 Dalton. The b-ions are calculated by the sum of monoisotopic masses of all composing amino acids plus one hydrogen monoisotopic mass while y ions are calculated as the sum of monoisotopic masses of all composing amino acids plus monoisotopic mass of one hydronium ion.

### Modified ResNet-18

The structure of modified ResNet-18 is shown in Figure S6. The main modifications include structural modifications and weight modifications. For structural modifications, the output size of the last layer in the modified ResNet-18 is modified to 2 to suit the binary classification problem. For weight modifications, the weights of the first 17 convolutional layers in the modified ResNet-18 are fine-tuned using liver cancer and thyroid cancer training data. The size of the subtensor is 224 x 224 x M, where the subtensor is the input of ResNet-18 and M is the number of precursor isolation windows. In the convolution and pooling operations, the dimension is generally reduced by 2, so the input size is preferably in the form of *b* × 2^*n*^. Furthermore, 224 x 224 is the size of the input layer commonly used in ResNet, facilitating us to directly call the pre-trained weights on the ImageNet dataset. Therefore, the input size 224 x 224 of the conventional ResNet is inherited in our model to extract the information embedded in the mass spectrum tensor.

### Deep neutral network implementation

For a binary classification problem, the final outputs represent the probabilities of the input being negative (normal samples) and positive (cancerous samples). A *softmax(·)* active function is used to map the output values to the range of [0, 1]. Then the cross entropy is used to defined the loss function.

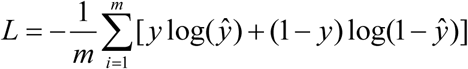

where m is the batch size, which is set to 8 in our model, *y* is the true label of the input, and *ŷ* is the the predicted score of the input being cancerous. During the training process, the model is iteratively trained 128 times on the dataset with Adam optimizer. The initial learning rate is set to 2×10^−3^, and after 64 iterations, the learning rate is changed to 2×10^−4^.

Lastly, the modified ResNet-18 was implemented in Tensorflow-1.14.0 and Python 3.7 running on GeForce GTX 2080 Ti GPUs.

